# GUN1 regulates tetrapyrrole biosynthesis

**DOI:** 10.1101/532036

**Authors:** Takayuki Shimizu, Nobuyoshi Mochizuki, Akira Nagatani, Satoru Watanabe, Tomohiro Shimada, Kan Tanaka, Yuuki Hayashi, Munehito Arai, Sylwia M. Kacprzak, Dario Leister, Haruko Okamoto, Matthew J. Terry, Tatsuru Masuda

## Abstract

The biogenesis of the photosynthetic apparatus in developing chloroplasts requires the assembly of proteins encoded on both nuclear and chloroplast genomes^1^. To co-ordinate this process there needs to be communication between these organelles, and while we have a good understanding of how the nucleus controls chloroplast development, how the chloroplast communicates with the nucleus at this time is still essentially unknown^2^. What we do know comes from pioneering work in which a series of *genomes uncoupled* (*gun*) mutants were identified that show elevated nuclear gene expression after chloroplast damage^3^. Of the six reported *gun* mutations, five are in tetrapyrrole biosynthesis proteins^4-6^ and this has led to the development of a model for chloroplast-to-nucleus retrograde signaling in which ferrochelatase 1 (FC1)-dependent heme synthesis generates a positive signal promoting expression of photosynthesis-related genes^6^. However, the molecular consequences of the strongest of the *gun* mutants, *gun1*^*7*^, is unknown, preventing the development of a unifying hypothesis for chloroplast-to-nucleus signaling. Here, we show that GUN1 directly binds to heme and other metal-porphyrins, affects flux through the tetrapyrrole biosynthesis pathway and can increase the chelatase activity of FC1. These results raise the possibility that the signaling role of GUN1 may be manifested through changes in tetrapyrrole metabolism and supports a role for tetrapyrroles as mediators of a single biogenic chloroplast-to-nucleus retrograde signaling pathway.

---

It has long been established that the expression of a large number of nuclear genes encoding chloroplast proteins is dependent on the presence of intact chloroplasts^8^ and mutations affecting chloroplast function or treatments with inhibitors such as norflurazon (NF) or lincomycin (Lin) result in the strong downregulation of many photosynthesis-related genes^7,9,10^. Mutants that lack this ability to co-ordinate the nuclear and chloroplast genomes were identified by their ability to retain expression of *LHCB1.2* after an inhibitory NF treatment^3^ and have been the basis for this research for the last 25 years. Of the original 5 *gun* mutants described, *gun2* and *gun3* lack a functional heme oxygenase 1 and phytochromobilin synthase^4^, *gun4* led to the identification of the Mg-chelatase regulator GUN4^5^, and *gun5* was mutated in the H subunit of Mg-chelatase (Supplementary Fig. 1)^4^. More recently, the identification of a dominant *gun6* mutant with increased FC1 activity^6^ led to the proposal that heme synthesized by FC1 is either the signal itself, a precursor of the signal or is required to generate the signal. However, very little progress has been made in further elucidating the signaling mechanism or in establishing whether this is the only biogenic retrograde signal. One barrier to tackling this problem is that the *gun1* mutant^7^, has been suggested to act independently from the tetrapyrrole-mediated GUN signaling pathway^4,11,12^. The GUN1 protein is a pentatricopeptide repeat (PPR) protein with a small MutS-related (SMR) domain^7^. The function of GUN1 is unknown, but it has been proposed to have a role in plastid protein homeostasis^13-16^, and can interact with proteins involved in both plastid protein synthesis and the tetrapyrrole pathway^13^

To further explore the interaction of GUN1 with tetrapyrrole biosynthesis, we first tested whether GUN1 could alter the flow through the tetrapyrrole pathway^17^ (Supplementary Fig. 1). Fig. 1a shows that feeding the precursor 5-aminolevulinic acid (ALA) to two *gun1* mutant alleles resulted in increased accumulation of protochlorophyllide (Pchlide) compared to wild-type (WT), while seedlings overexpressing GUN1^13^ had reduced accumulation of Pchlide (Fig. 1a). As ALA synthesis is the rate-limiting step in the pathway, one possibility to explain the altered Pchlide levels is that GUN1 is leading to a redistribution of tetrapyrrole to the heme branch. However, analysis of heme levels in dark-grown seedlings showed that *gun1-1* had more heme and GUN1ox lines had less heme than WT (Fig. 1b) indicating that GUN1 is impacting on flux through both branches of the pathway. Such a conclusion is consistent with the observation that *gun1* can rescue the reduction of heme caused by the *sig2* mutation^10^. Next, we tested whether overexpression of GUN1 also resulted in increased sensitivity to NF. As shown in Fig. 1c, both GUN1ox lines showed a stronger response to NF for *HEMA1* and *LHCB2.1* expression than WT seedlings, while, as expected, *gun1* rescued gene expression very strongly. Together these results show that altering GUN1 levels affects tetrapyrrole metabolism and that these changes correlate with changes in nuclear gene expression.

**Figure 1.**
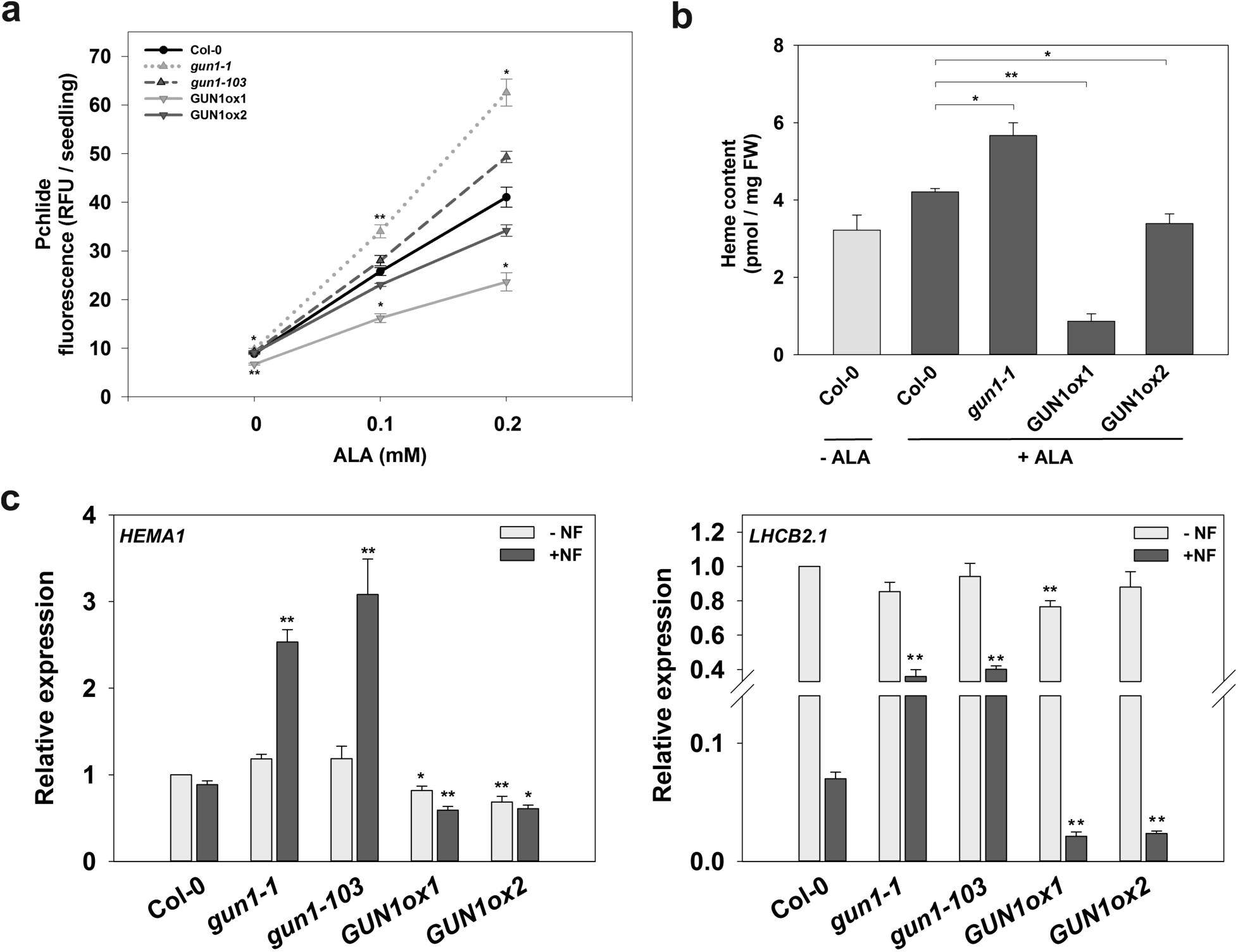
GUN1 affects tetrapyrrole metabolism. **a,** Protochlorophyllide accumulation in WT (Col-0), *gun1-1, gun1-103* mutants and GUN1ox1 and GUN1ox2 overexpressor lines grown 4 d in the dark on half-strength Murashige and Skoog medium supplemented with 1% (w/v) agar, 5 mM MES, (pH 5.8) and with or without 0.1-0.2 mM 5-aminolevulinic acid (ALA). 30 seedlings were analysed for each replicate. **b,** Total heme accumulation in seedlings treated with or without 0.2 mM ALA, as described in (a). **c,** Reverse transcription quantitative PCR (RT-qPCR) analysis of *HEMA1* and *LHCB2.1* transcript levels in WT (Col-0), *gun1-1, gun1-103,* GUN1ox1 and GUN1ox2 seedlings grown on half-strength Murashige and Skoog medium supplemented with 1% (w/v) agar with or without 1 µM NF under the following conditions: 2 d dark, 3 d WLc (100 µmol m^-2^ s^-1^). Expression is relative to Col-0 -NF and normalised to *YELLOW LEAF SPECIFIC GENE 8* (*YLS8*, At5g08290). Data shown are means +SEM or ± SEM of three independent biological replicates. Asterisks denote a significant difference vs. Col-0 for the same treatment, Student’s *t*-test (*p < 0.05; **p < 0.01).

Previously, it was shown by yeast two-hybrid analysis that GUN1 could interact with four tetrapyrrole biosynthesis enzymes including ferrochelatase 1 (FC1) (Supplementary Fig. 1)^13^. Although two FC isoforms are present in the Arabidopsis genome, GUN1 interaction was specific to FC1 and since an increase in FC1 activity is associated with increased nuclear gene expression in the *gun6* mutant^6^, we hypothesized that GUN1 may affect FC1 activity. To test this possibility, we attempted to express both GUN1 and FC1, but failed to express full-length GUN1 protein (Supplementary Fig. 2a, b). Prediction of the secondary and tertiary structure of GUN1 suggested a highly disordered domain in the N-terminal region (Supplementary Fig. 3), which may destabilize the GUN1 protein. Removal of this N-terminal domain allowed us to obtain GUN1 protein containing PPR and SMR motifs (GUN1-PS; Supplementary Fig. 2). We also expressed recombinant Arabidopsis FC1 as a glutathione-*S*-transferase (GST) fusion protein. The obtained GST-FC1 showed enzyme activity as measured by an increase in Zn-protoporphyrin (ZnProto) fluorescence (Fig. 2a)^18^. We then evaluated the effect of GUN1-PS on the ability of FC1 to catalyze Zn-chelation. The addition of GUN1-PS to FC1 enhanced the Zn-chelatase activity linearly with increasing concentration of GUN1-PS (Fig. 2a). In comparison, the same concentration of BSA had only a slight effect on activity, suggesting GUN1-PS did not merely stabilize FC1 (Fig. 2a). GUN1-PS itself had no Zn-chelating activity. Analysis of Michaelis-Menten kinetics revealed that addition of GUN1-PS decreased *K*_M_ values from 26.5 µM to 4.5 µM (Fig. 2b), suggesting GUN1-PS stimulated Zn-chelatase activity by enhancing substrate affinity for FC1.

**Fig. 2.**
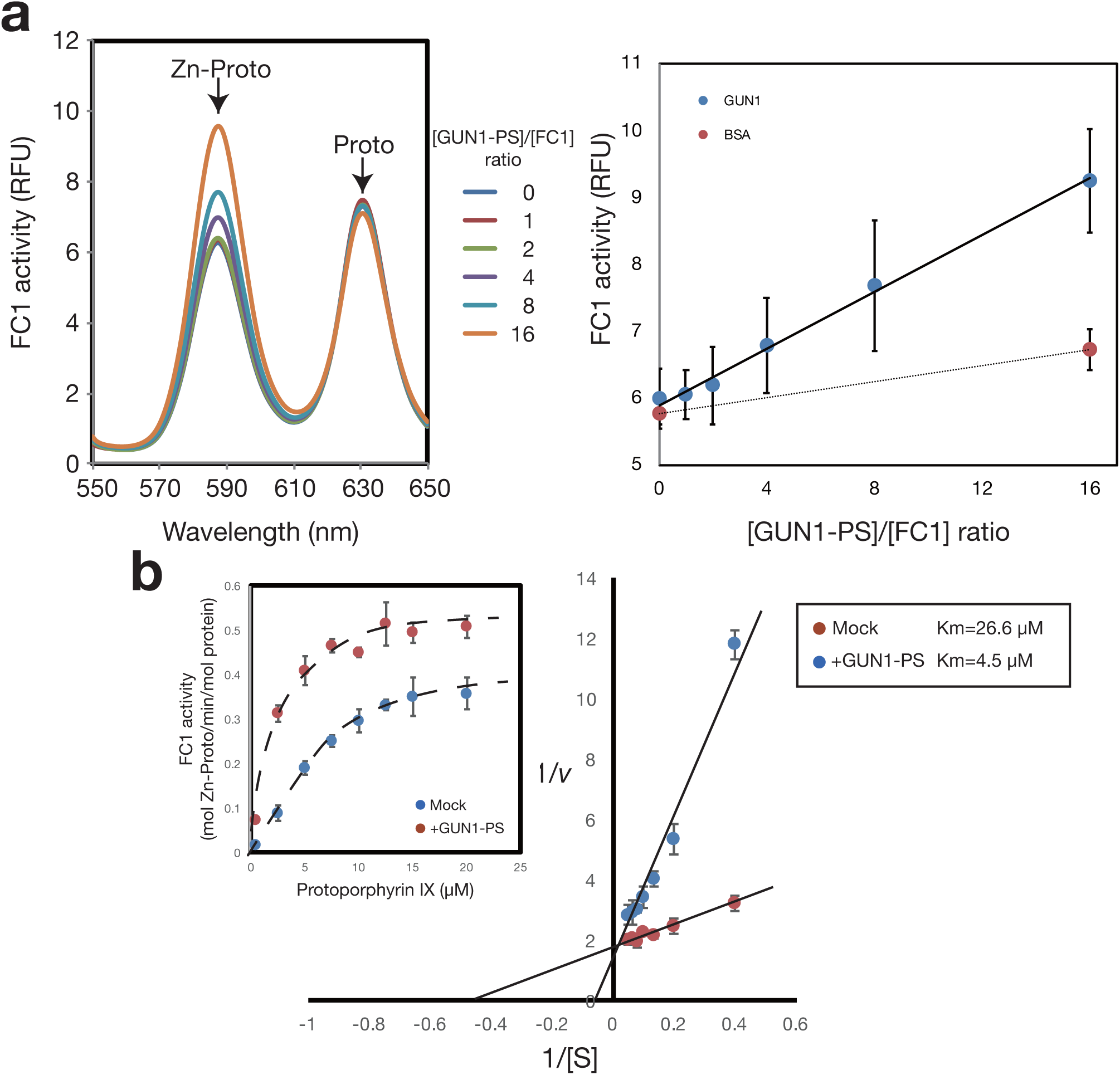
GUN1-PS enhances ferrochelatase 1 (FC1) activity. **a,** Arabidopsis FC1 protein expressed as a GST-fusion protein showed Zn-chelatase activity. Addition of GUN1-PS enhanced the formation of Zn-protoporphyrin IX (Zn-Proto) from Protoporphyrin IX (Proto) linearly with increasing concentration of GUN1-PS. Bovine serum albumin (BSA) was used as a negative control. **b**, Double reciprocal plot analysis of Zn-Proto formation by FC1 in the presence or absence of GUN1-PS. Inset shows Michaelis–Menten plot of the same data. *K*_M_ values of FC1 in the presence or absence of GUN1-PS are shown. Data shown are means +SEM (or ± SEM) of three independent replicates.

It has previously been shown that GUN4 can enhance Mg-chelatase activity by directly binding Mg-protoporphyrin (MgProto), the product of this reaction^5^. To examine whether a similar stimulating mechanism is employed in GUN1-dependent enhancement of FC1 activity, we tested the ability of GUN1-PS to bind heme using hemin-agarose beads. As shown in Fig. 3a, GUN1-PS demonstrated hemin-binding activity. To identify domains required for hemin binding, a series of truncations were constructed (Supplemental Fig. 2a) and tested for hemin binding (Fig. 3b). GUN1 proteins containing the PPR motifs (PPR1 and PPR2) showed significantly more hemin binding than those containing only the SMR motifs (SMR1 and SMR2). During the heme-binding assay, we observed that the colour of the hemin solution changed upon GUN1-PS binding (Fig. 3c inset), suggesting that changes in the hemin spectrum had occurred on binding. Spectrophotometric analysis showed that the GUN1-hemin complex exhibited a red shift of the Soret band and Q-band peaks compared to unbound hemin, consistent with specific binding (Fig. 3c). To determine the affinity of GUN1 for hemin, GUN1-PS was incubated with increasing hemin concentrations and binding determined by differential spectrophotometry (Fig. 3d). The increase in absorbance of the shifted Soret peak was plotted against porphyrin concentration (Fig. 3e) and a dissociation constant (*KD*) for the binding of GUN1-PS to hemin was estimated to be 6.08±1.11 µM using non-linear regression analysis and assuming a one-site binding model. This value sits within the range measured for a variety of heme-binding proteins^19^. Similar analyses for GUN1 binding to other metal-porphyrins resulted in estimated *KD* values for MgProto and ZnProto of 8.65±1.80 µM, and 3.10±0.86 µM, respectively (Supplementary Fig. 4). Thus, GUN1-PS has similar binding affinities for all three metal-porphyrins.

**Fig. 3.**
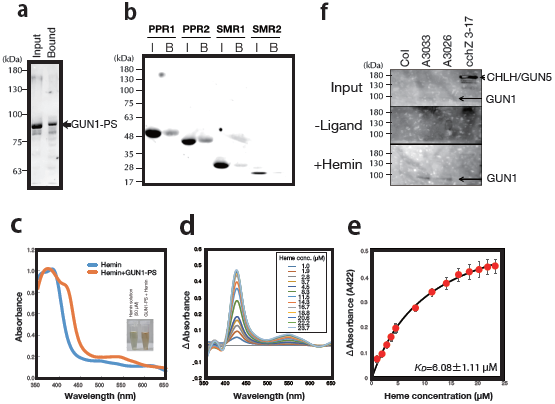
Recombinant GUN1 protein binds to heme through PPR motifs. **a**, Binding of GUN1-PS to hemin-agarose beads. **b**, Binding of a truncated series of GUN1 proteins (see Supplementary Fig. 2a) to hemin-agarose beads. I, input; B, bound. **c**, Absorption spectra of hemin and hemin-GUN1-PS complexes. Inset: photograph of hemin solution (50 µM) and hemin-GUN1-PS complex purified by gel filtration. **d**, Absorbance difference spectra of hemin-GUN1-PS minus hemin solution at different hemin concentrations. **e**, Change in absorbance of the Soret peak plotted against hemin concentration was used to determine the dissociation constant (*Kd*) of the heme-GUN1-PS complex assuming a one-site binding model. **f**, Binding of FLAG-tagged GUN1 isolated from Arabidopsis lines A3022 and A3026 (overexpressed in a *gun1* mutant background) to hemin-beads. The GUN5 protein (expressed in the *cchz* mutant background) is shown as a control.

Finally, to test whether GUN1 can bind to hemin in plant extracts, we constructed Arabidopsis lines expressing FLAG-tagged GUN1 under the control of its own promoter in the *gun1-102* mutant background (lines A3022 and A3026). As a control we also expressed FLAG-tagged GUN5 (line *cchZ* 3-17) in the *cchZ GUN5*-deficient mutant^20^. The *gun1*-*102* phenotype was complemented by GUN1 expression with de-repression of *LHCB1.2, RBCS1A* and *CHLH* expression by Lin rescued in these lines (Supplementary Fig. 5). Since GUN1 accumulates detectable levels only at very young stages of leaf development^16^, proteins were extracted from 4-day-old seedlings and subjected to the hemin binding assay (Fig. 3f). FLAG-tagged GUN1 was observed in the fraction bound to hemin, while the GUN5 protein, which also binds porphyrins^21^, was not. Thus, the binding of GUN1 to hemin shows specificity.

Our results suggest that GUN1 has two, likely independent functions, in modifying tetrapyrrole metabolism (Fig. 4). Firstly, it was able to restrict flow through both branches of the tetrapyrrole pathway such that feeding ALA resulted in reduced accumulation of both Pchlide and heme in dark-grown seedlings. The mechanism for this restriction is unknown but may be related to the observation here that GUN1 can bind metal-porphyrins and/or that it can interact with the tetrapyrrole enzymes porphobilinogen deaminase and uroporphyrinogen III decarboxylase 2 that catalyze shared steps in the tetrapyrrole pathway^13^. GUN1 is not a very abundant protein^16^ and it is unlikely that any restriction of the flow of tetrapyrroles is due to sequestration. Rather, we propose that the mode of action of GUN1 is regulatory. Moreover, the rapid degradation in the light^16^ would permit an increased flow of tetrapyrroles at a time when the demand for photosynthetic tetrapyrroles, both heme and Chl, is at its greatest.

**Fig. 4.**
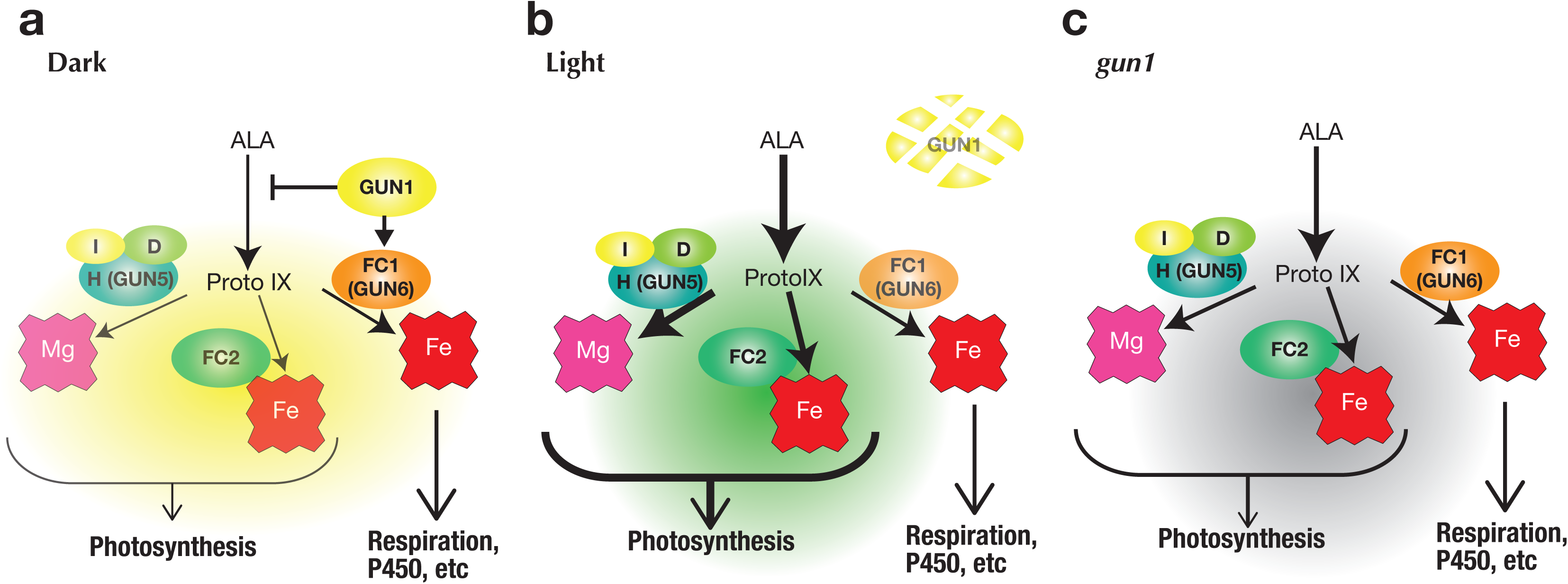
Model for GUN1 function in tetrapyrrole metabolism. **a,** In the dark GUN1 represses flow through the tetrapyrrole pathway, but promotes FC1 activity to ensure a sufficient supply of heme to the rest of the cell. **b**, In the light GUN1 is degraded promoting total tetrapyrrole synthesis required for chloroplast development. Under these conditions, FC1 activity is no longer promoted. The increased tetrapyrrole flux ensures a sufficient supply of substrate to FC1 to supply heme to the rest of the cell. **c**, In dark-grown *gun1* mutants tetrapyrrole synthesis is greater than in a wild-type seedlings and will continue to be during early stages of de-etiolation.

The second molecular function of GUN1 identified in this study, is the enhancement of FC1 activity through a more than 5-fold reduction in *K*_M_ (Fig. 2). It is likely that GUN1 stimulates FC1 activity through enhancing substrate affinity in a similar way to GUN4 enhancement of Mg-chelatase activity^5^. However, it should be noted that GUN1 product binding is more than 10-fold weaker than GUN4 (*K*_*D*_ value for Mg-deuteroprotoporphyrin binding to GUN4 is 0.26 µM^5^). The control protein used for this assay, BSA, also binds heme^22^, and thus enhancement of FC1 activity by GUN1 is quite specific. Nevertheless, this result was surprising given that GUN1ox lines showed reduced heme levels and it is probable that GUN1-dependent FC1 stimulation does not reflect the total heme content. We did not test whether GUN1 could also promote FC2 activity, but given that the interaction of GUN1 is specific for FC1^13^ this is not likely. To understand why GUN1 restricts tetrapyrrole synthesis, but promotes FC1 activity will take more detailed analysis of heme metabolism in young seedlings. However, one possible explanation is that GUN1 enhances FC1 activity to ensure a supply of heme to the rest of the cell under conditions in which tetrapyrrole synthesis is maintained at a low level (i.e. before the demand for photosynthetic tetrapyrroles). Once seedlings move to the light, the flow through the tetrapyrrole pathway is greatly increased through transcriptional upregulation of tetrapyrrole synthesis genes^23^. This would negate the need for GUN1 to promote FC1 to ensure a sufficient cellular heme supply and degradation of GUN1 would redress the balance between FC1 and FC2 to favour FC2 activity required to synthesize photosynthetic hemes^24^.

The observations presented here demonstrate that GUN1 alters tetrapyrrole metabolism (Fig. 4) and therefore that all described *gun* mutations affect this pathway. This observation therefore supports a model in which tetrapyrroles are mediators of a single biogenic retrograde signal during de-etiolation. The prevailing model is that FC1-dependent heme synthesis is required to generate a positive signal^6,25^. Our observations on the restriction of heme synthesis by GUN1 in seedlings are consistent with this model as it would be expected to result in more flow through FC1. However, the demonstration that GUN1 enhances FC1 activity does not appear to support it and may require the development of a new hypothesis. One scenario that could reconcile these two observations is if the restriction of tetrapyrrole synthesis by GUN1 was a more significant effect than enhancement of FC1, such that overall there was still less FC1-dependent heme in the presence of GUN1 than in its absence. Our own data showing that heme levels were reduced overall in GUN1ox lines do support such an interpretation. Alternatively, it is possible that binding of FC1-synthesized heme by GUN1 blocks release or propagation of the retrograde signal. In this case, GUN1 degradation in the light would ensure an increased signal to promote further chloroplast protein synthesis for continued development and the supply of new chloroplasts.

## Methods

### Plant material and growth conditions

*Arabidopsis thaliana* wild type (WT), the mutants *gun1-1, gun1-102*^7^, *gun1-103*^26^, and the GUN1ox lines^13^ were in the Columbia-0 (Col-0) ecotype. For physiological analyses, seeds were sown onto half-strength Murashige and Skoog (MS) medium (Melford Laboratories, Ipswich, UK) supplemented with 1% (w/v) agar (pH 5.8) and incubated in white light (WL; 120 µmol.m^-2^.s^-1^) for 2 h to induce germination. For Pchlide and heme assays seedlings were then grown for 4 d dark at 22 °C with or without 0.1 or 0.2 mM aminolevulinic acid (ALA; Sigma Aldrich) with 5mM MES. For gene expression analyses, seedlings were grown with or without 1 µM NF (a gift from John Gray, University of Cambridge, UK) in the absence of sucrose, kept for 2 d in the dark, followed by 3 d in continuous WL (WLc, 100 µmol.m^-2^.s^-1^) at 22 °C. For analysis of gene expression in complemented *gun1-102* lines, seeds were sown onto MS medium supplemented with 0.8% (w/v) agar (pH 5.8) and 2% (w/v) sucrose, with or without 560 µM Lin and grown for 4 d under WLc (100 µmol.m^-2^.s^-1^) at 23°C.

### Pchlide measurements

Pchlide levels were measured from 4-d old dark-grown seedlings. Seedlings were harvested into liquid nitrogen under green safe light, covered with foil and stored at − 80°C until required. Seedlings were ground under green safe light in a mortar and pestle on ice with 2 x 400 µL cold alkaline acetone solution (acetone: 0.1 M NH_4_OH; 9:1 (v/v)). Samples were vortexed briefly and centrifuged at 16,000 x *g* for 5 min at room temperature. Supernatants from two sequential extraction steps were collected and Pchlide levels were measured using a fluorescence spectrophotometer F-2000 (Hitachi High-Tech, Japan). Pchlide emission for the peak at ∼636 nm following excitation at 440 nm was recorded.

### Heme detection by chemiluminescence

Total non-covalently-bound heme measurements were performed following the protocol^27^ with minor modifications. 4-d old dark seedlings were harvested into liquid nitrogen under green safe light, covered with foil and stored at −80°C until required. Seedlings were ground in liquid nitrogen to a powder, extracted twice with 1 mL of acidic acetone (acetone:1.6 M HCl (80/20, v/v) and centrifuged at 16,000 x *g* at 4°C for 15 min to remove cell debris. Cleared extracts were diluted 1:1000 with sterile water. A 1 mM hemin stock solution (bovine hemin chloride, Sigma Aldrich, UK) was prepared in DMSO to create a calibration curve. To reconstitute horseradish peroxidase (HRP), 10 µL of extract or hemin standard was mixed with 40 µL of apo-HRP enzyme (BBI Solutions, Sittingbourne, UK) in 100 mM Tris-HCl, pH 8.4. The final enzyme concentration in each sample was 2.5 nM apo-HRP. Samples were vortexed, spun briefly and incubated for 30 min at room temperature. After transferring samples to a 96-well white polystyrene plate, 50 µL of HRP substrate (H_2_O_2_:luminol, 1:1; Immobilon^TM^ Western Chemiluminescent HRP Substrate, Merckmillipore) was added to each sample, mixed by pipetting and incubated for 5 min. Chemiluminescence was recorded using micro-plate reader (GloMax® 96 Microplate Luminometer, Promega) with 0.5 s integration time.

### Heme-binding assays

Different versions of GUN1 recombinant proteins (10 µM) in binding buffer (20 mM HEPES-NaOH (pH 7.9), 100 mM KCl, 1 mM MgCl_2_, 0.2 mM CaCl_2_, 10% (v/v) glycerol, 1 mM DTT, 0.2 mM PMSF, 0.1% Tween 20) were assessed for their ability to bind hemin-agarose beads (Sigma-Aldrich). For detection of GUN1-FLAG in Arabidopsis, total proteins were extracted from 4 d-old whole seedlings in the binding buffer. Hemin-immobilized FG beads^28^ prepared by Tamagawa Seiki (Tokyo, Japan) were used for assessment. Beads and protein were mixed for 2 h on a rotator at 4°C. Beads were extensively washed with the binding buffer, re-suspended in 2x Laemmli sample buffer containing 4% (v/v) β-ME, and heated for 5 min at 100°C before loading onto SDS gels. Proteins were transferred by western blotting to HyBond ECL membrane (GE Healthcare) and detected using anti-HisTag (MBL Life Science, Nagoya, Japan) and anti-DYKDDDDK (Fuji Film Wako) antibodies for the recombinant and *in planta* GUN1 proteins, respectively.

### Ferrochelatase assay

FC1 activity was measured by Zn-chelating activity as described previously^18^. The reaction mixture contained 10 µM protoporphyrin IX, 50 µM ZnSO_4_, and various ratios of GST-FC1 and GUN1-PS proteins in a total volume of 100 µL. The reaction was started by adding protoporphyrin IX solution, and was carried out at 30°C for 10 min, and then stopped by addition of 900 µL acetone. After centrifugation at 10,000 x *g* for 5 min, fluorescence emission spectra of the cleared supernatant were recorded between 550-650 nm with excitation at 420 nm in a FP-6200 fluorescence spectrometer (JASCO, Tokyo, Japan). Non-enzymatic formation of Zn-Proto formation was verified with denatured controls.

### Absorbance spectroscopy

Metal-porphyrin (hemin, MgProto, ZnProto) solutions were prepared by solubilizing in DMSO, followed by dilution in 20 mM Tris-HCl, pH 8.0. For assays, 10 µM GUN1-PS protein was subjected to UV-visible absorbance spectroscopy at room temperature in a Ultrospec 2100 pro spectrophotometer (GE Healthcare Bioscience) using 20 mM Tris-HCl buffer as a reference. Spectra were recorded between 350 and 650 nm using a 1 cm path length cuvette, with a metal porphyrin concentration range of 0–25 µM. Difference spectra were obtained by subtracting the metal-porphyrin spectrum from that of the protein-metal porphyrin complexes. Emerging peaks in the Soret region corresponding to metal-porphyrin-GUN1-PS complexes were plotted. Quantitative analysis of peak emergence was used to determine *K*D values for metal-porphyrin binding assuming one-site ligand binding model as described^29^. Titration data were analyzed using non-linear regression (OriginPro8 software; OriginPro Corporation, Massachusetts, USA).

### Circular dichroism (CD) spectrometry

CD spectra were measured in a JASCO J-805 spectropolarimeter (JASCO, Tokyo, Japan) at 200-250 nm with a quartz cuvette of 1 mm path length at 25°C (controlled by a thermostat circulating water bath). Analysis was performed using recombinant GUN1-PS proteins obtained during the serial dilution of urea at a protein concentration of 0.2 mg/mL.

### RNA extraction and gene expression analysis by RT-qPCR

RNA extraction and RT-qPCR analysis was performed as described previously^30^. Total RNA was extracted from around 100 mg of 5 d-old whole seedlings using a phenol:chloroform method, treated with RNase free DNase I (Promega Corporation, Madison, USA) and 1 µg RNA was used for cDNA synthesis with equal amounts of oligo(dT) and random nonamer primers using the nanoScript2 reverse transcription kit (Primerdesign Ltd., Southampton, UK). Transcript levels were measured by reverse transcription quantitative PCR (RT-qPCR) using Sybr Green master mix (Primerdesign Ltd.) and a StepOnePlusTM Real Time PCR System (Applied Biosystems, Foster City, USA). The thermal cycling conditions used were 95 °C for 10 min, followed by 40 cycles of 95 °C for 15 s and 60 °C for 60 s. Expression was normalized to *YELLOW LEAF SPECIFIC GENE 8* (*YLS8*, At5g08290). For analysis of gene expression in complemented *gun1-102* lines, total RNA was extracted from 4 d-old whole seedlings using the Agencourt Chloropure System (Beckman Coulter, Miami, USA) according to the manufacturer’s instructions. RNA was treated with RNase free DNase I (Sigma-Aldrich) and 1 µg RNA was used for cDNA synthesis with oligo(dT)12-18 using Transcriptor first-strand synthesis kit (Roche, Basel, Switzerland). Transcript levels were measured by qRT-PCR using LightCycler 480 SYBR Green I Master (Roche) and a LightCycler 96 (Roche). The thermal cycling conditions used were 95°C for 5 min, followed by 40 cycles of 95 °C for 5 s, 55 °C for 10 s, and 72 °C for 20 s. Expression was normalized to *TUBULIN BETACHAIN 2* (*TUB2*, At5g62690). All of the genes analyzed by RT-qPCR, with their accession numbers and corresponding primer sequences are listed in Supplementary Table S2.

### Construction of GUN1-FLAG lines

The endogenous promoter region of *GUN1* (ca. 1.5 kb) was amplified with P1 and P2 primers (see Supplementary Table 1) from Col genomic DNA using PrimeStarMax (Takara Bio, Kusatsu, Japan) and inserted in *Hin*dIII-*Xba*I digested pGWB511^31^ by the SLiCE method^32^ to give p511G1pro. *GUN1* cDNA was reverse transcribed using total RNA isolated from Col with Transcriptor High Fidelity (Sigma-Aldrich, Tokyo, Japan). *GUN1* cDNA (ca. 2.8 kb) was amplified using G1f and G1r primers, and cloned into pENTR-D-TOPO vector to give pENTR-D-GUN1 (Thermo Fisher, Tokyo, Japan). p511G1pro was digested with *XhoI,* purified, and subjected to the Gateway LR reaction with pENTR-D-GUN1 to give p511G1pro-GUN1. p511G1pro-GUN1 was introduced to *Agrobacterium* strain GV3101(MP90) and used for transformation of *gun1-102*. The expression of GUN1-FLAG was confirmed by qPCR and western blotting using an anti-DYKDDDDK antibody (Fuji Film Wako, Osaka, Japan).

### Plasmid constructions and expression of recombinant proteins

For *in vitro* translation and transcription experiment, full-length *GUN1* cDNA (At2g31400) and pIVEX1.3 WG (Roche) fragments were amplified by PCR and ligated using the In-Fusion HD Cloning Kit (Takara Bio). Using RTS 100 Wheat Germ CECF Kit (Roche), GUN1 protein was expressed in RTS ProteoMaster (Roche). Full-length *GUN1* cDNA and N-terminal truncated versions as *Eco*RI-*Sal*I PCR fragments were ligated into pET48b (Novagen, Wisconsin, USA) and transformed into *E. coli* Rosetta gami2 (DE3) (Novagen). For other truncated versions of GUN1 (PPR1, PPR2, SMR1 and SMR2), primers were designed to exclude domains of interests, and PCR fragments were self-ligated (see Supplementary Table 1 for all primers). After pre-culture in LB medium containing 100 µg mL^-1^ kanamycin, GUN1-PS was induced by adding 0.1 mM IPTG at 37°C for 3 h. After collecting cells by centrifugation at 4,000 x *g* for 5 min, the cell pellet was re-suspended in 20 mM Tris-HCl, pH 8.0, and subjected to ultrasonic disruption. Since GUN1-PS formed inclusion bodies, the insoluble fraction was repeatedly washed with buffer containing 20 mM Tris-HCl (pH 8.0), 0.5 M NaCl, 1 M urea, 2% (v/v) Triton X-100, 20 mM DTT and 10 mM EDTA. The resultant insoluble fraction was dissolved in 20 mM Tris-HCl (pH 8.0) and 8 M urea, and subjected to dialysis with serial dilution of urea (6, 4, 2, 0 M urea). Full-length *FC1* cDNA was ligated into pGEX-2T vector (GE Healthcare, Buckinghamshire, UK) using the In-Fusion HD Cloning Kit (Takara Bio) and transformed into *E. coli* DH5*α*. After pre-culture, GST-FC1 fusion protein was induced by adding 0.1 mM IPTG at 16°C overnight. After collecting cells by centrifugation, the cell pellet was re-suspended in 20 mM Tris-HCl, pH 8.0, and 10% (v/v) glycerol and subjected to ultrasonic disruption. After centrifugation at 10,000 x *g* for 10 min, the supernatant was used directly in the ferrochelatase assay.

## Acknowledgement

We would like to thank Takehito Kobayashi and Takahiro Nakamura (Kyushu University, Japan) for bioinformatic analysis of PPR domain of GUN1. We thank Saumya Awasthi for contributions to pilot studies. We also thank Biomaterial Analysis Center, Technical Department of Tokyo Institute of Technology for technical support. We are grateful for funding from JSPS KAKENHI grant numbers JP16K07393, JP18H03941, JP16H02217, JP18K05386 and JP17K07444. M.A. was supported by the Institute for Fermentation, Osaka. S.M.K. was supported by the Gatsby Charitable Foundation. Work on retrograde signalling by M.J.T. is supported by the UK Biotechnology and Biological Sciences Research Council. D.L. is supported by the Deutsche Forschungsgemeinschaft (SFB-TR 175, project C05).

## Author Contributions

T.S. expressed recombinant proteins and conducted enzyme assays, analysed and interpreted data and contributed to writing the article. N.M. performed experiments, analysed and interpreted data and contributed to writing the article. A.N. analysed and interpreted data. W.S. and T.S. performed heme binding experiments. K.T. analysed and interpreted data. Y.S. contributed biophysical analysis including CD spectrometry. M.A. analysed and interpreted data. S.M.K. performed the physiological and gene expression analyses, analysed and interpreted data and contributed to writing the article. D.L. provided research materials and contributed to writing the article. H.O. analysed and interpreted data and contributed to writing the article. M.J.T. jointly conceived the study, analysed and interpreted data and co-wrote the article. T.M. jointly conceived the study, performed experiments, analysed and interpreted data and co-wrote the article.

---

**Supplementary Fig. 1.**
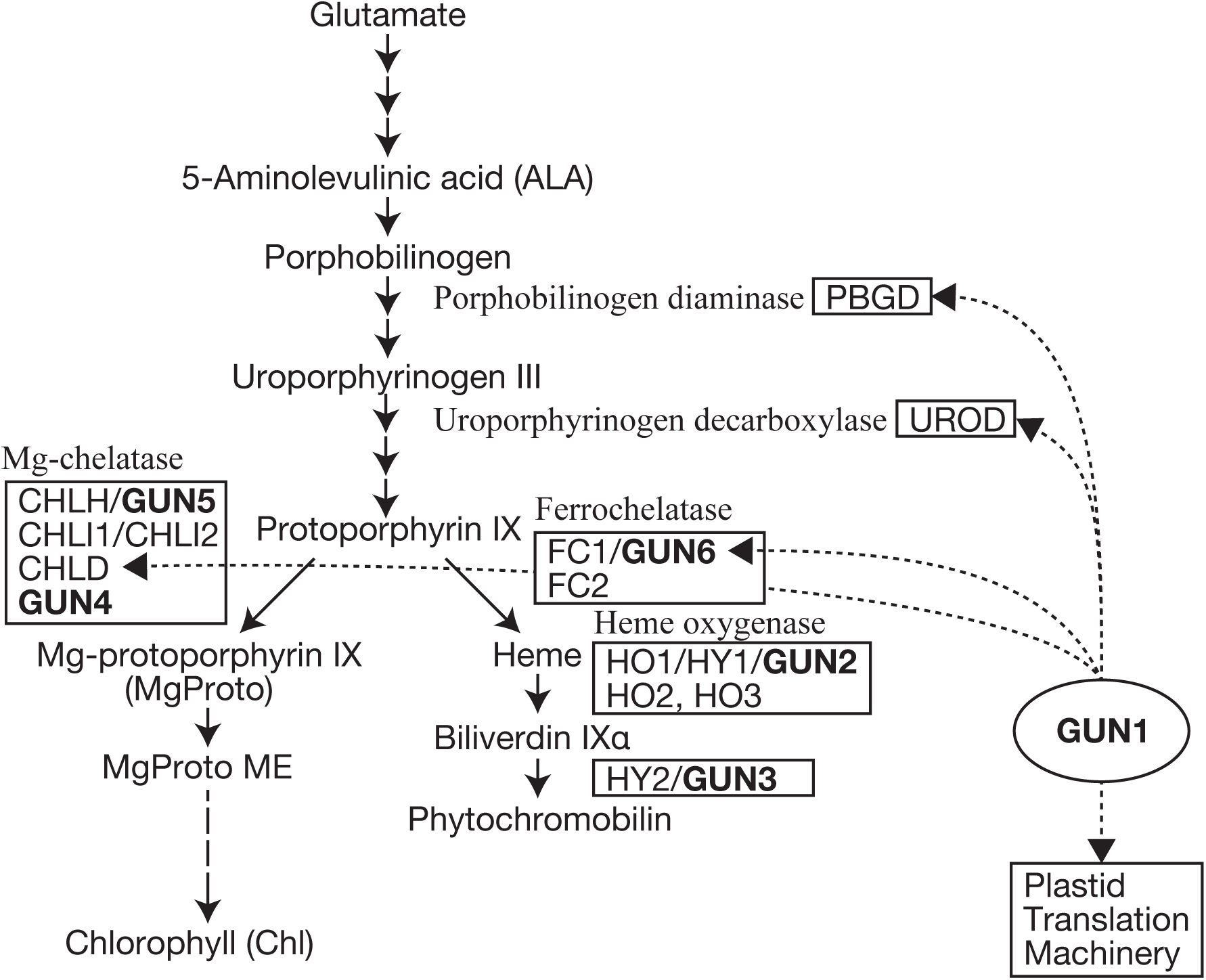
Tetrapyrrole biosynthetic pathway. A simplified schematic of tetrapyrrole biosynthesis in higher plants. Arrows indicate enzymatic steps; the proteins and genes involved are indicated to the right of the arrow. Boxed proteins and genes indicate the steps that result in a *gun* phenotype when blocked. Dashed lines indicate proteins that showed interaction with GUN1 in a yeast 2-hybrid analysis (Tadini et al., 2016).

**Supplementary Fig. 2.**
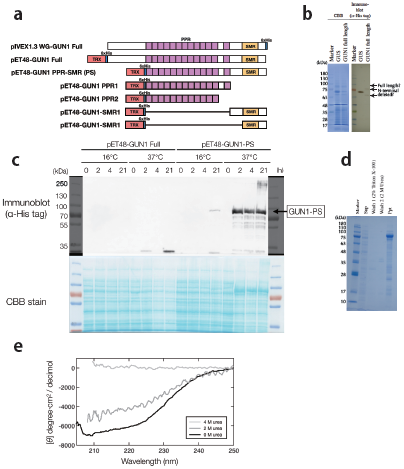
**a,** Schematic diagrams of recombinant GUN1 proteins used. **b**, Attempted expression of full-length GUN1 protein using *in vitro* transcription and translation system with the pIVEX1.3. **c**, Expression of full-length and N-terminal truncated GUN1 proteins in *E. coli*. Expression of recombinant GUN1 proteins were induced at 16°C and 37°C for 0, 2, 4 and 21 h. **d**, Purification of GUN1-PS inclusion body. GUN1-PS was solubilized in buffer containing 8 M urea and refolded by serial dilution. **e**, CD spectra of GUN1-PS during serial dilution of the urea concentration. Note that initial attempts to express recombinant full-length GUN1 protein by *in vitro* transcription and translation and in *E. coli* (**a**) were unsuccessful (**b, c**). Therefore, we truncated the N-terminal region and successfully expressed the GUN1 protein containing the PPR and SMR motifs as an N-terminal thioredoxin (Trx)-His_6_-fused protein (**a**). This recombinant protein was expressed as an inclusion body, which required treatment with 8 M urea to denature the protein, and subsequent serial dilution to isolate a refolded recombinant protein. In this fraction, GUN1-PS comprised more than 95% of the total protein (**d**) and CD analysis suggested that substantial refolding of GUN1-PS had occurred (**e**).

**Supplementary Fig. 3.**
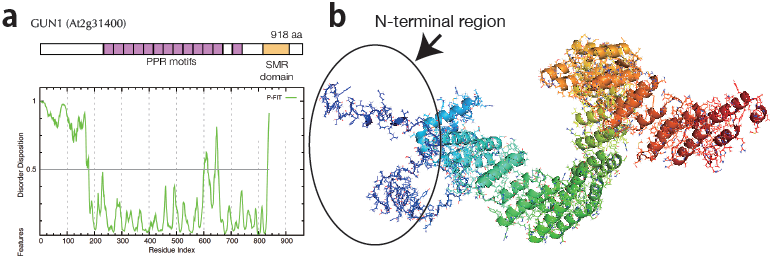
Prediction of secondary and tertiary structures of Arabidopsis GUN1. **a,** Schematic diagram of GUN1 and prediction of the disordered region of the GUN1 protein (PONDR-FIT of DisProt (http://disorder.compbio.iupui.edu/)). **b**, Predicted 3D structure of GUN1 by I-TASSER (https://zhanglab.ccmb.med.umich.edu).

**Supplementary Fig. 4.**
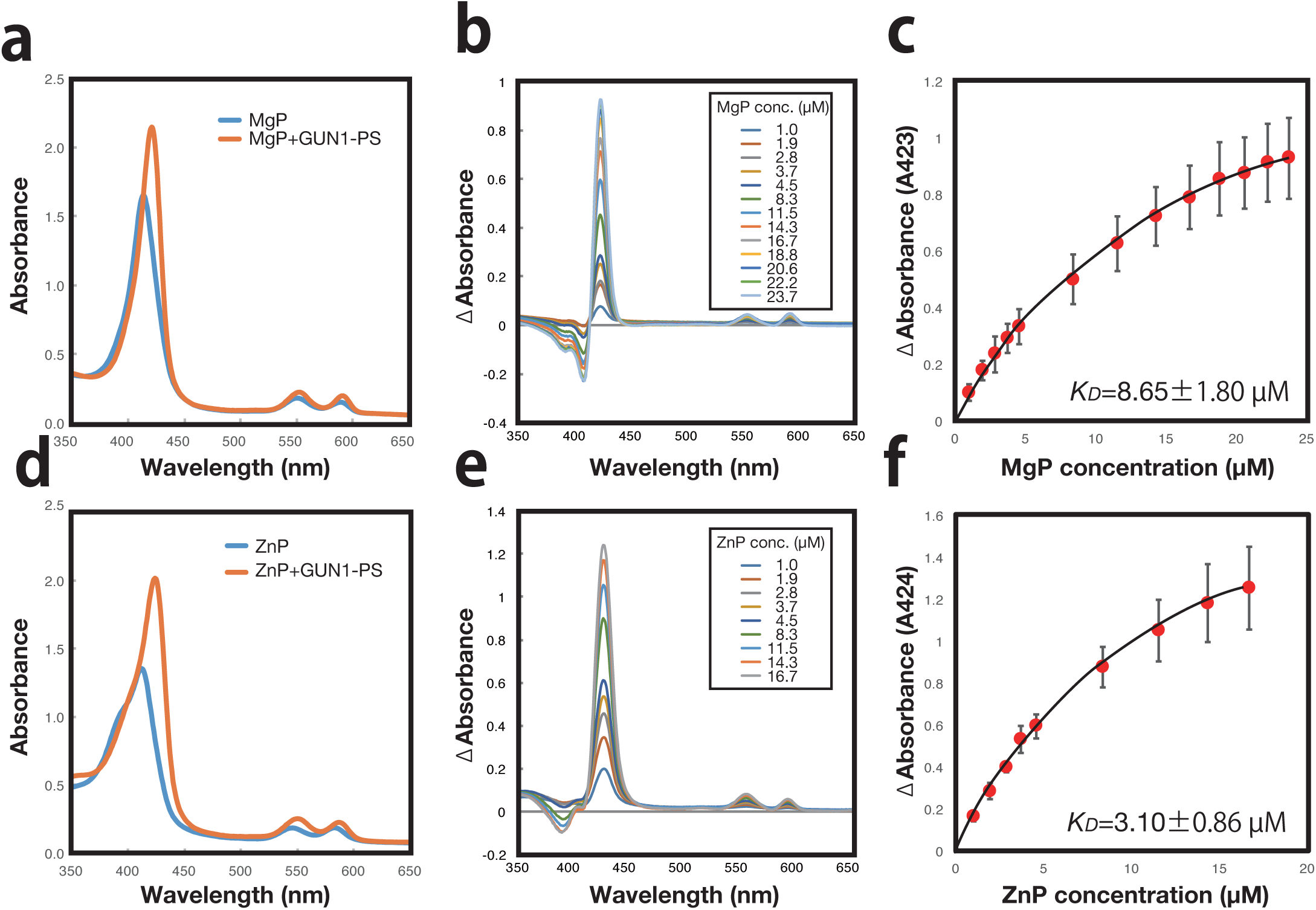
Spectral changes of metal-porphyrin-GUN1-PS complexes. **a, d,** Absorbance spectra of metal-porphyrins (Mg-protoporphyrin IX (MgP), and Zn-protoporphyrin IX (ZnP)) and metal-porphyrin-GUN1-PS complexes, respectively. **b, e**, Absorbance difference spectra of MgP-GUN1-PS (b) or ZnP-GUN1-PS (e) minus metal porphyrin solution at different porphyrin concentrations. **c, f,** Change in absorbance the Soret peak plotted against MgP (c) or ZnP (f) concentration was used to determine the dissociation constant (*K*D) of the metal porphyrin-GUN1-PS complexes assuming a one-site binding model.

**Supplementary Fig. 5.**
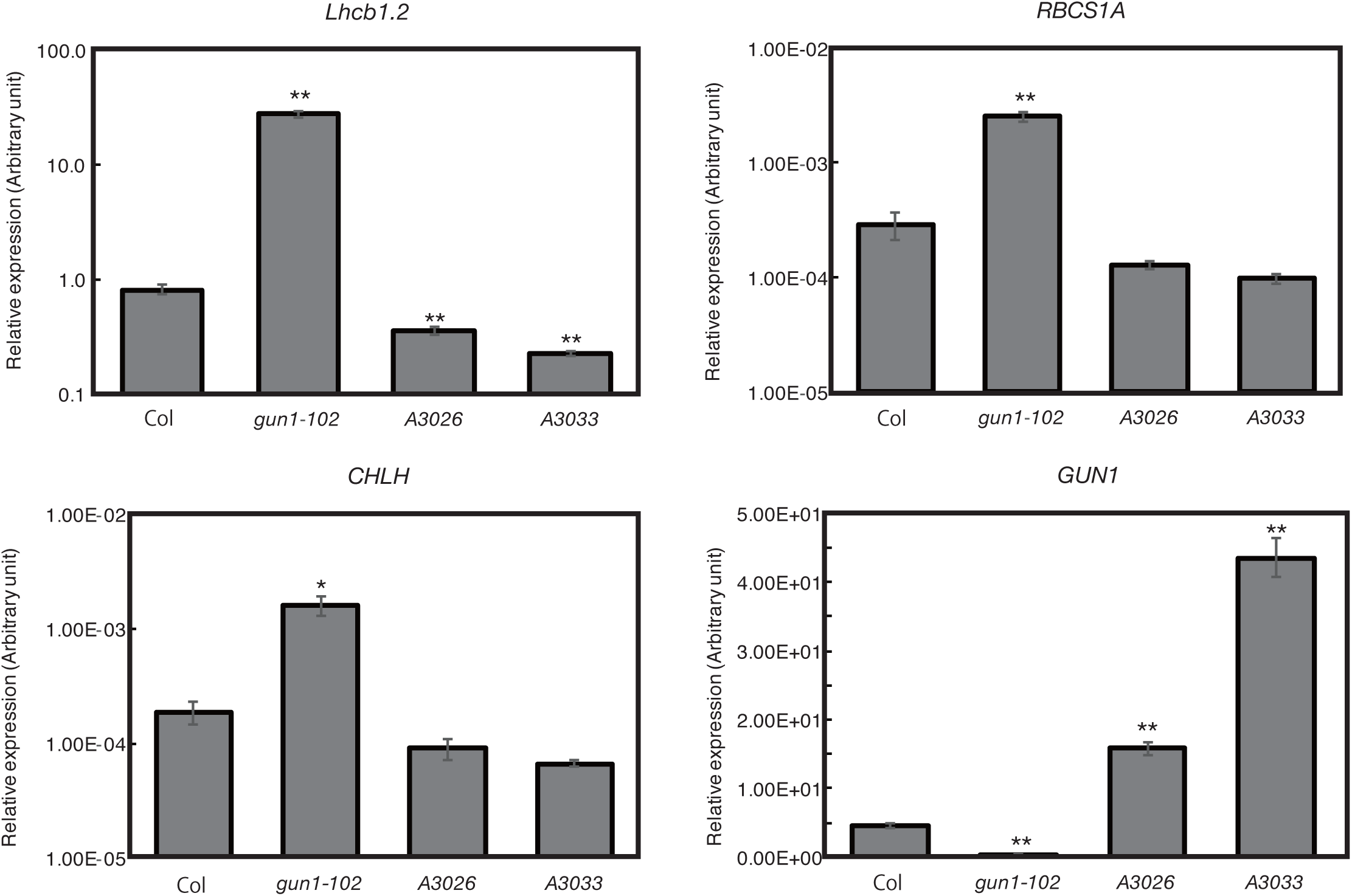
Complementation of *gun1-102* with *GUN1pro::GUN1-FLAG*. Seedlings were grown on Murashige and Skoog medium supplemented with 2% (w/v) sucrose and 0.8% (w/v) agar (pH 5.8), with or without 450 µM Lin for 4 d in continuous WL (100 µmol m^-2^ s^-1^). Expression was determined by RT-qPCR and is normalized to *TUBULIN BETACHAIN 2* (*TUB2*, At5g62690). Genotypes correspond to wild-type (Col), *gun1-102*, and complementation lines (A3026 and A3033) expressing *GUN1pro::GUN1*-*FLAG*. Data shown are the means ±SEM of three independent biological replicates. Asterisks denote a significant difference vs. Col-0 for the same treatment, Student’s *t*-test (*p < 0.05; **p < 0.01).

**Supplemental Table 1.**
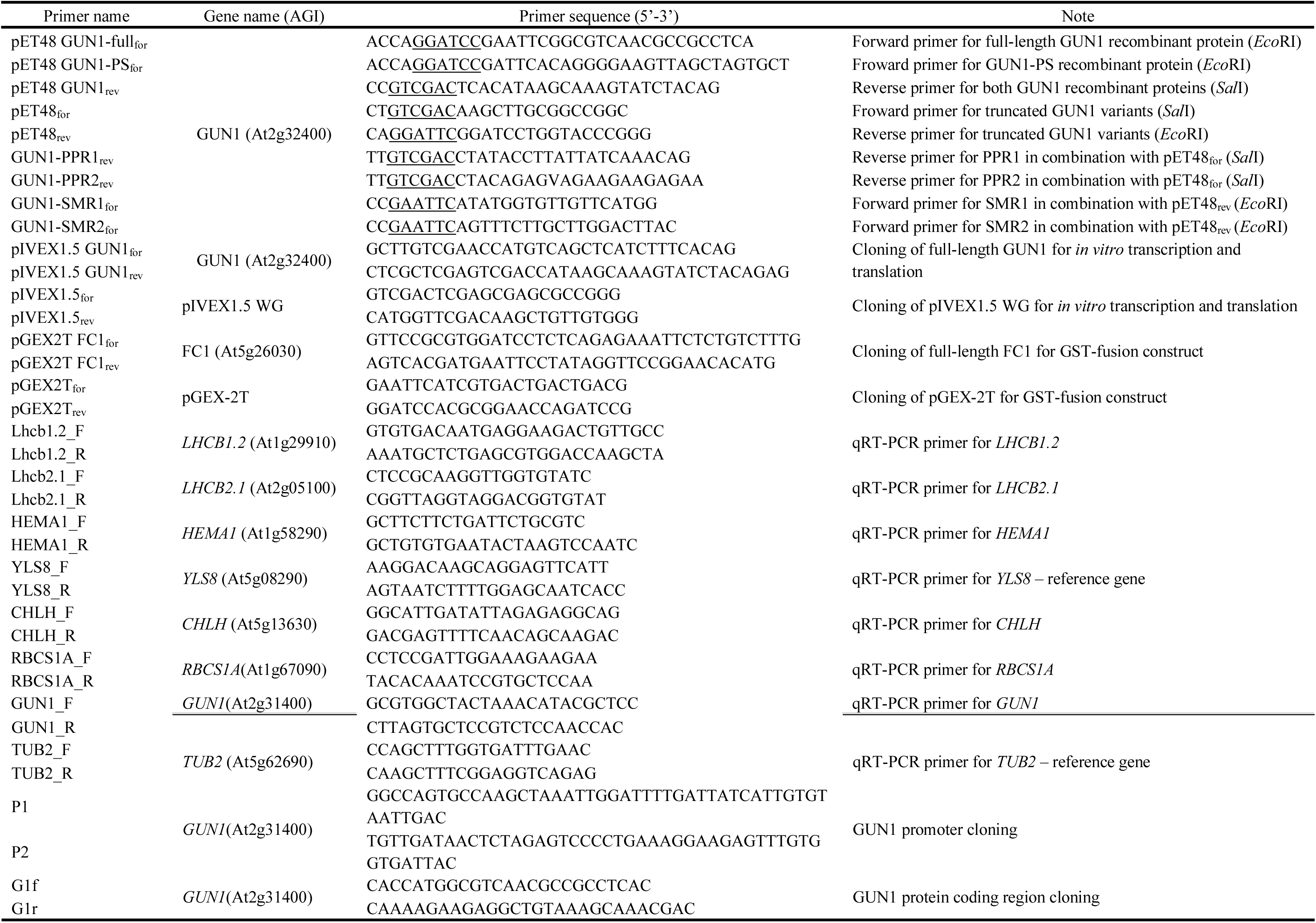
Primer sequences used in this study

